# Cryptic diversity and population structure at small scales: The freshwater snail *Ancylus* (Planorbidae, Pulmonata) in the Montseny mountain range

**DOI:** 10.1101/054551

**Authors:** Jan N. Macher, Martina Weiss, Arne J. Beermann, Florian Leese

**Affiliations:** Aquatic Ecosystem Research, University of Duisburg-Essen, Universitätsstraße 5, 45141 Essen, Germany

**Keywords:** cryptic species complex, population structure, barcoding, species distribution modelling, freshwater invertebrates

## Abstract

Anthropogenic impacts like intensified land use and climate change are severe threats to freshwater biodiversity and effective biodiversity monitoring is therefore one of the most urgent tasks. This is however often hampered by the lack of knowledge regarding the number and ecology of species. Molecular tools have shown many freshwater taxa to comprise morphologically cryptic species, which often occur in sympatry on a small geographic scale. Here, we studied the freshwater snail *Ancylus fluviatilis* (MUELLER, 1774) species complex in the Iberian Montseny Mountains. We hypothesised 1) that several species of *A. fluviatilis* sensu lato occur in the Montseny, 2) that different *Ancylus* species seldom co-occur in syntopy due to different ecological demands or interspecific competition, and 3) that species show a pattern of strong population structure within streams or catchments due to ecological preferences or local adaptation. We barcoded 180 specimens from 36 sites in the Montseny for the cytochrome c oxidase subunit I (COI) barcoding gene and molecularly identified two *Ancylus* species. These species seldom occurred in syntopy and a species distribution modelling approach showed differing bioclimatic preferences of the species. One species mainly occurs in cooler, higher altitude streams while the second species occurs in lower-altitude areas with higher temperatures. Tests of population structure showed that both species possibly do not disperse well in the study area and that populations within species are likely adapted to certain bioclimatic conditions in different regions of the Montseny. Our results highlight the need to incorporate molecular techniques into routine monitoring programmes.

## INTRODUCTION

Anthropogenic impacts like intensified land use and climate change are severe threats to biodiversity (Vörösmarty et al., 2010; Steffen et al., 2015). Therefore, monitoring the ongoing loss of biodiversity is highly important and many countries worldwide haveestablished programmes to do so. However, the total loss of biodiversity can only be monitored when accurate knowledge on the number, ecology, distribution and genetic diversity of species is available. For many areas and ecosystems, such information does often either not exist or is inaccurate: in recent years, the use of molecular methods has shown that the numberof species is underestimated in many taxa (e.g. Amato et al., 2007; Pfenninger and Schwenk, 2007; Adams et al., 2014). This is especially true for freshwater ecosystems, which harbour a large number of morphologically indistinguishable or cryptic animal species (e.g. Pauls et al., 2010; Weigand et al., 2011; Weiss et al., 2014). The ecology of most cryptic species is however rarely known since it has been studied for relatively few taxa only (e.g. Ortells et al., 2003; Rissler and Apodaca, 2007; Lagrue et al., 2014; Fišer et al., 2015). This lack of knowledge poses a risk, since monitoring programmes and biodiversity assessments can come to wrong conclusions if species with different ecologies are treated as being identical regarding their ecological demands and thus, their suitability to indicate ecosystem health (e.g. Macher et al., 2016). Also, extinction events and loss of biodiversity can go unnoticed.In this regard, using molecular methods to study freshwater species in mountain ranges is especially promising since many mountain ranges have been shown to harbour a large number of cryptic freshwater species (e.g. Pauls et al., 2009; Katouzian et al., 2016; Mamos et al., 2016). Further, mountain ranges comprise many different habitats due to their topographic and climatic complexity, often leading to different species communities occurring within a small geographic area(Finn and Leroy, 2005; Múrria et al., 2014; Cauvy-Fraunié et al., 2015). Topography and climate form natural barriers to dispersal and many taxa occurring in mountain ranges show phenotypic and genetic adaptation to the highly differing conditions along the altitudinal gradient (Liebherr, 1986; Bonin et al.,2006; Keller et al., 2013; Watanabe et al., 2014), ultimately leading to mountain ranges being centres of high biodiversity. Using molecular methods to study species in such environments can help understand species diversity, ecology and their possible genetic adaptation to different habitats. Further, it can allow inferring the potential loss of species and genetic diversity when the environment changes.

Here, we analysed diversity and spatial distribution patterns in a common European stream invertebrate taxon, the freshwater limpet *Ancylus fluviatilis* (MÜLLER, 1774) sensu lato, in the Montseny mountain range on the Iberian Peninsula. The Montseny is partof the Catalan pre-coastal range (North East Iberian Peninsula). It is located at the intersection of the warm and arid climate of the mediterranean lowlands and the cooler and more precipitation rich climate of the mountainous region reaching to the Pyrenees (Thuiller et al., 2003). There are three main catchments within this area, all of which are characterised by steep altitudinal gradients ranging from less than 500 to 1706 metres above sea level (masl) within approximately 10 kilometres and thus, comprising highly variable climatic conditions (Peñuelas and Boada, 2003; Jump et al., 2007). We chose to study the widespread hololimnic freshwater limpet *Ancylus fluviatilis* sensu lato, because itisknown to comprise several cryptic species (Hubendick, 1970; Pfenninger et al., 2003; Albrecht et al., 2006). Of those species, Clade 1 and Clade 4 (Pfenninger et al., 2003) potentially co-occur in the North East Iberian Peninsula but have never been foundin the here studied area. On a European scale, Pfenninger et al. (2003) found that the different *A.fluviatilis* species differ significantly in their ecological demands:while Clade 1 prefers cooler areas with precipitation-rich summers, Clade 4 occurs mainly in arid, generally hotter areas. Both climatic conditions can be found in the Montseny,making it an ideal area for studying the number and distribution of species within the *A.fluviatilis* species complex. *A. fluviatilis* sensu stricto isable todisperse over longer distances (Cordellier and Pfenninger, 2008), e.g. by passive transport via waterbirds and other organisms (Rees, 1965), a phenomenon commonly found in other snails and freshwater molluscs (Rees, 1965; Boag, 1986; Van Leeuwen et al., 2012).Thecurrentdistribution of species in the *A.fluviatilis* species complex isthusexpectedto be limited by ecological demands (Cordellier and Pfenninger, 2008).

In this study, we expected 1) to find morphologically cryptic species of the *A. fluviatilis* species complex in the Montseny mountain range, 2) that different *A. fluviatilis* species seldom co-occur in syntopy due to different ecological demands or interspecific competition, and 3) that species show a pattern of strong population structure within streams or catchments of the Montseny due to ecological preferences or localadaptation to bioclimatic conditions in different altitude zones. In contrast, a weaker population structure is expected between streams and catchments located in the same altitude zone, because bioclimatic conditions are more similar here and *A. fluviatilis*isknown to be able to disperse over catchment boundaries, e.g. via vectors such as waterbirds.

To test these hypotheses, we first analysed the mitochondrial cytochrome c oxidase subunit 1 gene (COI) to determine the number and distribution of *Ancylus* species found in the Montseny. Second, we used a modelling approach based on bioclimatic variables to identify variables that might help to explain the occurrence of species and third, we performed population genetic analyses to investigate intraspecific partitioning of variation.

## MATERIALS AND METHODS

### Sampling

Sampling was performed in the Montseny mountain range (located on the North East Iberian Peninsula, Fig. 1a) and the direct surrounding area in September 2013. 44 sites were checked for the presence of *A. fluviatilis*, which was found in 36 of these sites (see Table A1 for coordinates). The three main catchments (Tordera, Besòs, Ter) and an altitudinal gradient from 120 masl to 1295 masl (Fig. 1b) were covered by the sampling. *Ancylus* specimens were collected by hand picking specimens from stones in the streams. All specimens were immediately stored in 70% ethanol,later transferred to 96% ethanol and stored at 4°C untilfurther analysis.

**Fig. 1.**
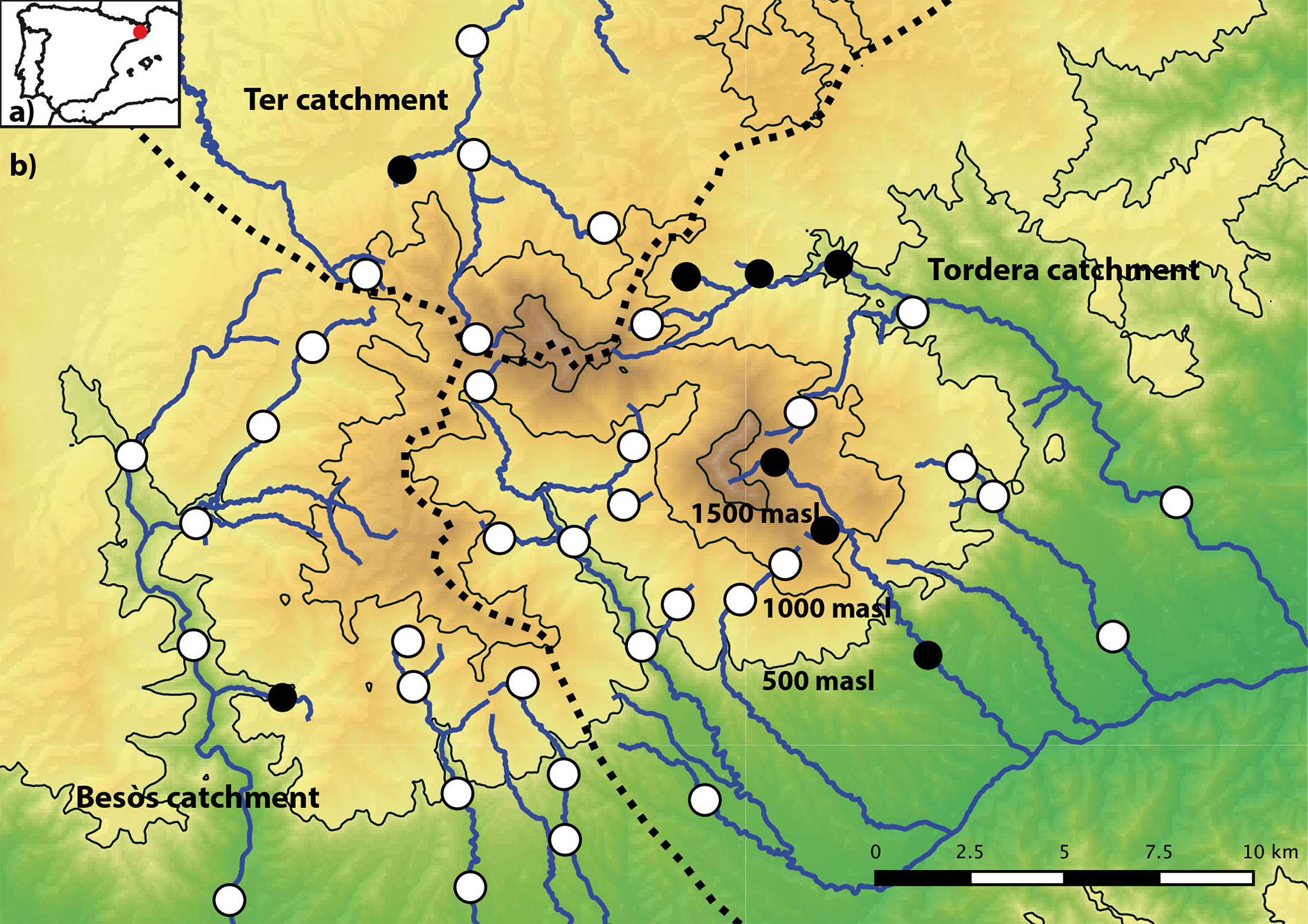
Figure 1: a) Location of the Montseny mountain range on the Iberian Peninsula (red circle); b) Location of the sampling sites in the Montseny. White dots: Sites where *Ancylus fluviatilis* sensu lato was found. Black dots: Sites where *A. fluviatilis* sensu lato was not found. Catchment boundaries are shown as dashed black lines. Rivers are shown as blue lines.

A sampling permit for protected areas (Parc Natural del Montseny) was obtained from the park management prior to sampling.

### DNA extraction, amplification, and sequencing

DNA was extracted from muscle tissue of 180 specimens (5 per site, 36 sampling sites) using a salt extraction protocol (Sunnucks and Hales, 1996) (Overview of the samples: Table A1).A 658 bp-fragment of the barcoding gene COI was amplified using the primers LCO1490 and HCO2198 (Folmer et al.,1994). The PCR mix was prepared using the following protocol: 1 x PCR buffer, 0.2 mM dNTPs, 1 μl of DNA template, 0.025 ᴜ/μl Hotmaster Taq (5 PRIME GmbH, Hilden, Germany) and 0.5μM of each primer. The mix was filled up to 25 μl with sterile H2O and placed in a thermocycler for amplification. PCR settings for the COI amplification were: initial denaturation at94°C for 2 min; 36 cycles of denaturation at 94°C for 20 s, annealing at 46°C for 30 s, extension at 65°C for 60 s; final extension at 65°C for 5 min. 9 μl of the PCR product were purified enzymatically with 10 ᴜ of Exonuclease I and 1 ᴜ Shrimp Alkaline Phosphatase (Thermo Fisher Scientific, Waltham) by incubating at 37°C for 25 min and a denaturation step at 80°C for 15 min. Bidirectional sequencing was performed on an ABI 3730 sequencer by GATC Biotech (Constance, Germany).

### Species delimitation

Raw reads were assembled and edited using Geneious 6.0.5 (Biomatters). The MAFFT plugin (v. 7.017,Katoh and Standley, 2013) in Geneious was used to compute a multiple sequence alignment (automatic algorithm selection, default settings). The final length of the cropped alignments was 655 bp. The alignment was translated into amino acids using translation table 5 (invertebrate mitochondrial codon usage table) to make sure that no stop codons were present.The best model of evolution for further analyses of the data was selected with jModeltest 2.1.2 (Darriba et al., 2012)(default settings). Fabox (Villesen, 2007) was used to collapse sequences into haplotypes. PopART (v.1, Leigh and Bryant, 2015) was used to create statistical parsimony haplotype networks (Clement et al., 2000) with a 95% connection limit.

Two approaches were used to test for the presence of cryptic species in *Ancylus fluviatilis* sensu lato: first, the tree-based Generalized Mixed Yule Coalescent (GMYC) approach (Pons et al., 2006) and second, the automated distance-based barcode gap determination approach (ABGD, Puillandre et al., 2012). An ultrametric tree for all unique COI haplotypes was calculated for the GMYC analyses using BEAST v.1.8.0 (Drummond et al., 2012). BEAST was run for 10 million MCMC generations, sampling every 100th tree and using both standard coalescent and the GTR+G sequence evolution model. Tracer v.1.6 (Rambaut et al., 2013) was used to test for effective sampling size(ESS) and convergence of parameters. TreeAnnotator v.1.8 (Rambaut and Drummond, 2013) was used to generate a linearized consensus tree, discarding the first 3000 trees as burn-in. R v.3.1.1 (R Core Team 2014) was used for analysis of the resulting tree with ‘SPLITS’ (Species Limit by Threshold Statistics) (Ezard et al., 2009) with the single threshold model to test for the presence of multiple species within the dataset. The second approach used for species delimitation was ABGD. Default settings were used, with Pmax=0.1 and the K2P-model of distance correction (Kimura, 1980), as this is the common approach in DNA barcoding studies. Once the number of genetic clades within the dataset was determined, specimens from each group were blasted against the Barcode of Life database (Ratnasingham and Hebert, 2007) to verify species assignment. The *Ancylus* sequences from Pfenninger et al. (2003, accession numbers AY350509 - AY350525) were downloaded and aligned with the sequences generated in this study to verify assignment of sequences to one ofthe known cryptic species. Alignments for each species were created with Geneious and networks were computed with popArt as described above. QGIS (v 2.8, available from www.qgis.org) was used to create distribution maps.

### Bioclimatic variables analyses

The bioclimatic preferences of species were modelled using a maximum entropy method in MaxEnt 3.3.3e (Phillips and Dudík, 2008), which has been shown to work well with small sample sizes (Pearson et al., 2007). The region modelled was part of the North East Iberian Peninsula (area between coordinates 42°C18’N, 1°C48’E, 41°C06’N, 3°C00’E; WGS84). Atotal of 19 climate layers in the 30 arc-seconds grid were obtained from WorldClim (Hijmans et al., 2005) and resampled to a cell size of 800 x 800 m. WorldClim datasets are based upon standard meteorological precipitation and temperature measurements, which are transformed into bioclimatic variables (Hijmans et al., 2005). These datasets are commonly used as predictor variables in species distribution modelling. To avoid using highly nonindependent variables in the analyses and thus omit overfitting of models, a Spearman’s rank correlation tests was performed across all pairs of variables using ENMtools (Warren et al., 2010) and R (R Core Team 2015). The correlation coefficient values used as thresholds beyond which values were treated as independent were 0.7, 0.8 and 0.9. All species presence points were used to build the model; 25% of the presence points were retained for training the model. All models were run 10 times with random partitioning of training and validation points. The accuracy of all computed models was evaluated with the area under receiver operation characteristic curve which was also used to choose models for use in further analyses (Boubli and De, 2009). Range overlap and bioclimatic niche overlap were computed by using Schoener’s D statistics as implemented in ENMtools. The values range from 0 (meaning no bioclimatic niche overlap) to 1 (identical range and bioclimatic niche, respectively).

### Geographic partitioning of genetic variation

Φ_ST_ values as an indicator of population subdivision were calculated separately for all species found in the *A. fluviatilis* species complex using the software Arlequin (v. 3.11, Excoffier et al., 2005). Φ_ST_ was chosen since it takes population history(number of mutations between haplotypes) into account. For analyses of population differentiation between altitude zones, populations of all species were classed in three groups: <500 masl, 500-1000 masl and >1000 masl as in Múrria et al. (2014). For analyses of population differentiation between catchments, populations were classed as belonging to one of the three catchments (Tordera, Besòs, Ter) and Φ_ST_values between groups were calculated.The Bayesian Clustering software GENELAND (v.4.0.5 as implemented in R; Guillot et al., 2005), was used to further analyse population structure in the found *Ancylus* species. Fabox was used to extract variable sites from alignments of the found species and PGDSpider (Excoffier and Lischer, 2010) was used to convert these alignments files into the GENELAND format. The settings used for running GENELAND were: Five independent runs with a maximum of 10 populations, 300 nuclei, 10 million iterations, thinning interval of 10 000, resulting in 1000 retained trees. The first 200 trees were discarded as burn-in.

## RESULTS

### Molecular species delimitation

*Ancylus* was found in 36 out of 44 sampled sites (Fig. 1b, see Table A1 for coordinates). A total of 180 specimens were analysed for the COI barcoding gene. The 655 bp alignment had 54 (9.2%) variable sites and a GC content of 29.6%. The null model of a single species was rejected both with the GMYC (likelihood ratio for single threshold model: 31.68, p<0.001) and the ABGD approach (Pmax 0.1%). Both ABGD and GMYC suggested the presence of two groups in *A. fluviatilis* sensu lato. Blast searches against the Barcode of Life database assigned all sequences of both molecularly identified clades to either *A. fluviatilis* Clade 1 or *A. fluviatilis* Clade 4, both submitted by Pfenninger et al. (2003). Alignment of the generated sequences with those obtained from Genbank clustered 102 specimens with *A. fluviatilis*Clade 1, while 78 sequences clustered with *A. fluviatilis* Clade 4. Both clades were defined by Pfenninger et al. (2003). Clade 1 corresponds to *A.fluviatilis* sensu stricto, while Clade 4 is a yet undescribed species with circum-mediterranean distribution (Pfenninger et al. 2003). The species are referred to as *Ancylus* C1 and *Ancylus* C4.

### Bioclimatic characterisation

For the modelling approach based on the 19 bioclimatic variables obtained from WorldClim,3, 6 and 10 variables were retained after Spearman’s Rank Correlation tests with thresholds of 0.7, 0.8 and 0.9, respectively. The 6 variable model with a Spearman’s RankCorrelationthreshold of 0.8 resulted in good area under the receiver operating characteristic curve (AUC) values for both species (*Ancylus* C1: 0.96, *Ancylus* C4: 0.89), thus this model was chosen for all further analyses to mediate between lower variable correlation and higher model fitting (see Table A2 for all AUC values and variables). The best explaining bioclimatic variables for the occurrence of *Ancylus* C1 were the variables bio7 (“Temperature Annual Range”) and bio19 (“Precipitation of Coldest Quarter”). Occurrence of *Ancylus* C4 was best predicted by the variables bio7 (“Temperature Annual Range”) and bio15 (“Precipitation Seasonality”) (Table A2). Bioclimatic niche overlap for *Ancylus* C1 and *Ancylus* C4 was 0.643, the range overlap computed for occurrence likelihoods of >50% was 0.728 (Table A3).

### Geographic partitioning of genetic variation within Ancylus species

Both *Ancylus* C1 and *Ancylus* C4 were found in all three studied catchments of the Montseny. *Ancylus* C1 was found at 22 sampling sites and *Ancylus* C4 at 17 sampling sites. Both species occurred in syntopy at4 sampling sites (11.43%; <500 masl zone: 1 site, 500-1000 masl zone: 2 sites,>1000 masl zone: 1 site; Fig. 2). *Ancylus* C1 was found more often at higher altitude sites (332 - 1295 masl, median 665 masl) than *Ancylus* C4, which was mainly found at lower altitude sites (120-1172 masl, median 440 masl) *Ancylus* C1 showed significant population differentiation between the Tordera and BesÒs (Φ_st_: 0.322, p=0.00001) and the Tordera and Ter catchment (Φ_st_ 0.167, p=0.0001)(Table A4). The most common haplotype (HC1_1) was found at 14 sites and in all three catchments (Tordera: 5 sites, Besòs: 4 sites, Ter: 5 sites)(Fig. 3a). HC1_2 was found at five sites, of which four are located in the Tordera catchment and one in the Ter catchment. HC1_3 was found at 9 sites and all three catchments (Tordera: 6 sites, Besòs: 2 sites, Ter: 1 site). Haplotypes C1_4, C1_5, C1_6, C1_7 and C1_8 were found in a maximum of two specimens each and at single sampling sites only. Significant population differentiation in *Ancylus* C1 was also found between the altitude zones <500 masl and >1000 masl (Φ_st_: 0.343, p=0.00001) and between 500-1000 masl and >1000 masl (Φ_st_: 0.274, p=0.0001)(Table A4). GENELAND found three geographically defined groups in *Ancylus* C1. Group 1 contains the sampling sites dominated by HC1_2, mainly lying above 1000masl (4 out of 5 sites). Group 2 contains sampling sites mainly dominated by HC1_1 (Populations in all altitude zones, but mainly (9 sites) in the 500- 1000 masl zone). Group 3 contains sampling sites mainly dominated by HC1_3 (500-1000 masl zone: 5 sites; <500masl zones: 3 sites) (Fig. 3a). A maximum of three substitutions were found between haplotypes of *Ancylus* C1 (Fig. 3b). *Ancylus* C4 showed significant population differentiation between the Tordera and Besos (Φ_st_: 0.408, p=0.0001) and the Besòs and Ter catchments (Φ_st_: 0.876, p=0.00001), respectively. The most common haplotype (HC4_3) was found at 13 sampling sites (BesÒs catchment: 9 sites, Tordera catchment: 4 sites). The second most common haplotype (HC4_1) was found in all three catchments (Tordera catchment: 4 sites, Ter catchment: 1 sites, BesÒs catchment: 1 site). The haplotypes C4_2, C4_4 and C4_5 were found at single sites and in single specimens only (Fig. 3c). In *Ancylus* C4, significant population differentiation was found between populations in the <500 masl and the 500-1000 masl zone (Φ_ST_: 0.335, p=0.009) as well as between the <500 masl and >1000 masl (Φ_ST_: 0.667, p=0.00001) zone. GENELAND found two geographically defined groups in *Ancylus* C4. Group1contains four sampling sites located in the northern and eastern parts of the study area, mainly dominated by haplotype C4_1 (>1000 masl zone: 2 sites, <500 masl zone: 2 sites). Group 2 contains 13 sampling sites in the mid and western part of the Montseny, mainly dominated by haplotype C4_3 (<500 masl zone: 10 sites, 500-1000 masl zone: 3 sites)(Fig. 3c). A maximum of two substitutions were found between haplotypes of *Ancylus* C4 (Fig. 3d).

**Figure. 2.**
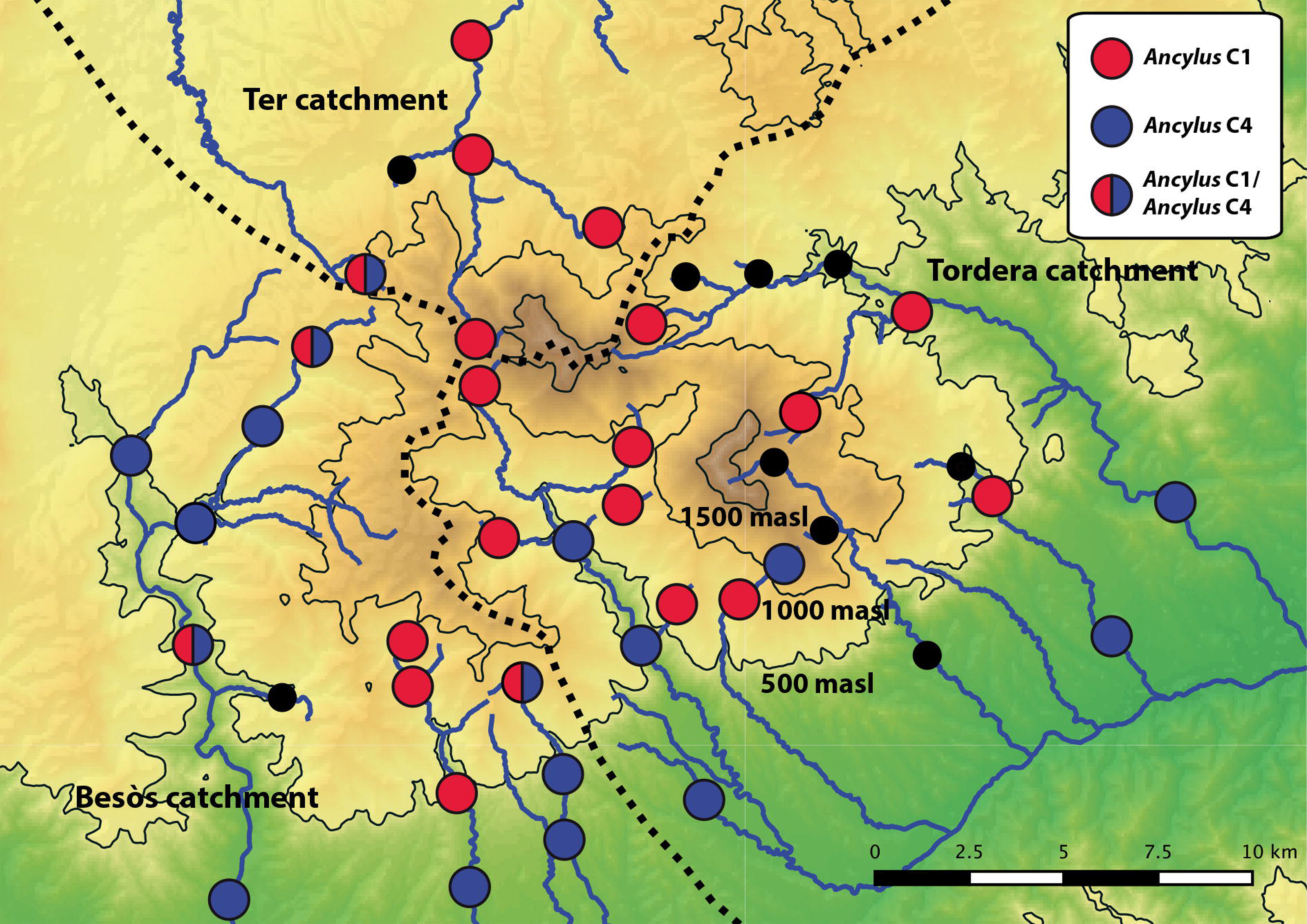
Map showing the occurrence of *Ancylus* Clade 1 and Clade 4 in the Montseny. Red dots indicate presence of *Ancylus* C1, blue dots presence of*Ancylus* C4. Mixed blue and red dots indicate the presence of both species at one sampling site. Black dots indicate absence of *Ancylus.* Catchment boundaries are shown as dashed black lines. Rivers are shown as blue lines.

**Figure. 3:a)+ c).**
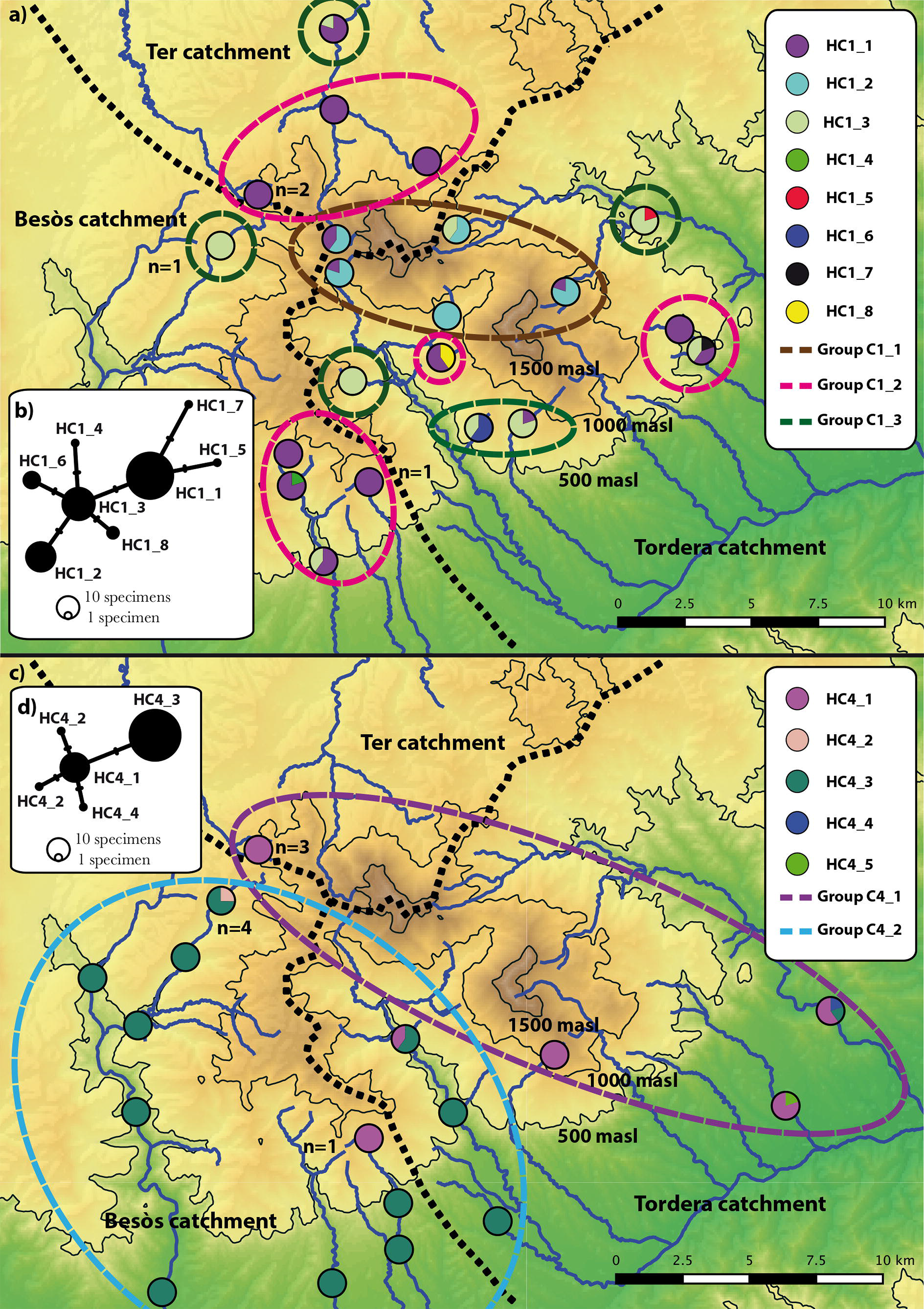
Map of the study area indicating the presence of *Ancylus* Clade 1 and *Ancylus* Clade 4, respectively, the number of studied specimens per sampling site and found COI haplotypes. Number of specimens per sampling sites is 5 unless otherwise stated. GENELAND groups are shown as coloured dashed lines and catchment boundaries as black dashed lines. b)+ d) Statistical Parsimony Network of *Ancylus* Clade 1 and *Ancylus* Clade 4 haplotypes, respectively. Dots represent sampled haplotypes, bars represent number of substitutions between haplotypes.

## DISCUSSION

In this study, we investigated the number, distribution and genetic variation of *Ancylus fluviatilis* sensu lato species in the Montseny mountain range on the Iberian Peninsula. Our first expectation was that cryptic species of *Ancylus fluviatilis* sensu lato are present in the study area. This expectation was met by the discovery of two species occurring in the Montseny. Both *Ancylus* species were initially delimited by Pfenninger et al. (2003). While*Ancylus* C1 corresponds to *A. fluviatilis* sensu stricto, *Ancylus* C4 is a yet undescribed species with a Mediterranean distribution, ranging from Portugal through the Southern Iberian Peninsula to Italy (Pfenninger et al., 2003; Albrecht et al., 2006).Although the distribution and ecology of cryptic *A. fluviatilis* sensu lato species are roughly known on a European scale, our study is the first that allows assessing the differences regarding small scale distribution, population structure and bioclimatic preferences of two *A. fluviatilis* sensu lato species in the same region, allowing for a better understanding of species’ ecologies. In the future, this might help with identification of species and improving stream quality assessments by making it possible to assign correct ecological traits to species.

Our second expectation was that different *A. fluviatilis* sensu lato species rarely occur in syntopy, possibly due to different ecological demands or competition. This expectation was clearly met, as *Ancylus* C1 and *Ancylus* C4 occurred in syntopy in only 11.43% of the sites and showed strong altitudinal partitioning: *Ancylus* C1 was found mainly at higher altitudes, while *Ancylus* C4 was mainly found at lower elevations. As elevation is a good indicator for bioclimatic and environmental conditions such as temperature (-0.65°C per 100m increase in altitude on average; Dodson and Marks, 1997) and flow velocity (due to steeper mountain slopes at higher altitudes), the observed pattern hints at different bioclimatic preferences of the two species. This is supported by the MaxEnt modelling approach, which suggests that *Ancylus* C1 occurs in areas with lower mean annual temperature and strong precipitation during the cooler season of the year, while *Ancylus* C4 occurs in areas with strong precipitation seasonality and higher mean annual temperature. These findings correspond to those of Pfenninger et al. (2003) who, on a European scale, found *Ancylus* C1 to mainly inhabit cooler, precipitation rich areas and *Ancylus* C4 to mainly inhabit hotter, seasonally dry areas. Similar patterns have been observed in other aquatic invertebrate species (e.g. Monaghan et al., 2005; Múrria et al., 2014). Closely related species co-occurring in the same area often inhabit different habitats due to different bioclimatic preferences or competition exclusion principle (e.g. Fišer et al., 2015). In southern Europe, cooler and precipitation rich conditions are mainly found at higher elevations, suggesting that *Ancylus* C1 might be close to the southern border of its distribution range in the Montseny. For aquatic species, higher precipitation means that more water is available during at least parts of the year, which is especially important in generally dry areas or areas with high precipitation seasonality.It appears thus possible that *Ancylus* C1 is more relying on constant flow of streams it inhabits, while *Ancylus* C4 might be able to cope with intermittent conditions in streams and generally higher water temperatures in lowland streams. This is especially important in the light of future climate change and ongoing human activities such as water abstraction, which might greatly alter temperature and precipitation patterns,ultimately changing flow regimes and thus possibly driving some species into local extinction while giving other species the possibility to colonise new habitats. Knowingthe number andecology of species in an area allows tracking the impact of such changes and preventing the loss of species.

Third, we expected the studied species to show a pattern of intraspecific altitudinal population structure due to local adaptation or ecological preferences, but not between catchments due to the reported dispersal abilities of *A. fluviatilis* and the fact that streams in the same altitude zone have equal bioclimatic conditions irrespective of the catchment they are located in. Tests of genetic differentiation, albeit based on a limited number of specimens and on the mitochondrial COI gene only, revealed that both *Ancylus* species found in the Montseny show a division into lower altitude and higher altitude populations that differ genetically. *Ancylus* C1 populations from above1000 masl significantly differed from populations below 1000 masl, which was also affirmedbythe GENELAND results. *Ancylus* C4 populations from below 500 masl differed significantly from populations located above that altitude, likely due to the fact that the common haplotype C4_3 was found mainly below 500 masl. GENELAND, however, did not confirm the existence of a haplotype group corresponding with altitude in *Ancylus* C4, probably due to the low number of specimens from above 500 masl sampling sites. A pattern of genetic differentiation between higher-altitude and lower-altitude populations could hint at two phenomena, possibly in combination: One possibility is that *A. fluviatilis* sensu lato species are weak dispersers that rarely migrate over longer distances within and between streams, thus over time populations diverge genetically due to limited geneflow.Weak dispersal capabilities have been found in other freshwater gastropods (Kappes and Haase,2012). However, at least *A. fluviatilis* sensu stricto has been shown to disperse over longer distances (Cordellier and Pfenninger, 2008), e.g. via passive transport (Rees, 1965). The second explanation could be that populations from higher and lower altitudes differ genetically due to selective processes, having adapted to the different bioclimatic conditions. This corresponds to findings in other studies that found high-altitude populations of species to be genetically different and potentially adapted to harsher bioclimatic conditions (e.g. McCulloch et al., 2009; Dussex et al., 2016). Fast adaptation to bioclimatic conditions has been found in *Ancylus* C1 (*A. fluviatilis* sensu stricto)(Cordellier and Pfenninger, 2008), possibly making this explanation for the pattern observed in the Montseny more likely. GENELAND analyses hint at *Ancylus* C1 being a stronger disperser than *Ancylus* C4, as the common haplotype C1_1 was evenly found in all three catchments, while a clear East West division of haplotypes was found in *Ancylus* C4. Dispersal through waterbirds or mammals, as has been found in snails and other freshwater taxa (Segerstråle, 1954; Figuerola and Green, 2002; Haun et al., 2012; Van Leeuwen et al., 2012), might be more common in *Ancylus* C1 than in *Ancylus* C4, possibly indicating that the latter species is a weaker disperser. It can only be speculated that this might be due to different dispersal strategies of the species, different preferred microhabitats that make it unlikely for *Ancylus* C4 to attach to waterbirds or due to lower survival rates when being out of the water. Also, it is possible that *Ancylus* C4 is not good at establishing new populations when arriving in habitats already occupied by*Ancylus* C1, potentially due to lower competitiveness. Further studies addressing the ecologies of both species need to be conducted, ideally using a combination of laboratory and field experiments. Significant population differentiation in *Ancylus* C1 between catchments in the Montseny can be explained by the fact that no population containing the haplotype dominating above 1000m (HC1_1) is located in the Besoscatchment andonly one is found in the Ter catchment. Population differentiation in *Ancylus* C4 is high between the Tordera and Besòs catchment, which was also confirmed by the GENELAND results. An East West segregation of haplotypes exists in C4, largely corresponding to the catchments Tordera and Besòs and possibly hinting at different colonisation events in the past or adaptations to bioclimatic conditions, which could not be identified in this study. The high population differentiation between Ter and Besos catchments should beneglected due to only one sampling site from the Ter catchment being included in the analyses. Overall, it remains possible that the pattern observed is mainly due to generally low genetic variation within the species and the relatively low number of sequenced specimens. The data is based on a limited number of specimens only and needs to be interpreted with care. Studies involving nuclear markers and possibly a greater number of specimens per site are needed to verify the observed patterns of genetic variation on a geographically small scale. However, the patterns of population differentiation between altitudinal zones and catchments demonstrate the need to take population structure into account when planning to protect species and ecosystems. Environmental changes in the lower or higher altitude zones or in catchments might lead to the extinction of adapted genotypes. The resulting lower levels of genetic diversity possibly limit the adaptive potential of species and their abilityto respondto environmental changes, ultimately leading to loss of genetic diversity in the species as a whole and increasing the risk of extinction (Bálint et al.,2011).

Our study highlights the importance and potential of using molecular techniques to study species diversity and genetic diversity of species. Molecular studies can greatly help to understand the impact of climate change and other human stressors on biodiversity (Bálint et al., 2011; Hampe and Jump, 2011; Pauls et al., 2013; Macher et al., 2016). Here, we found cryptic species within a common freshwater taxon and genetic divergence within species on a small geographic scale. Our results show that patterns of genetic diversity, connectivity and bioclimatic preferences can be different even between closely related species, a fact that should be considered in biomonitoring and conservation plans. Knowledge of freshwater species’ diversity and ecological preferences is also important due to the fact that many bioassessment and monitoring programs worldwide rely on species occurrence data as a metric to measure ecosystem quality (e.g. Carter and Resh, 2001; Stark et al., 2001; Haase et al., 2004) and the indication value of species is mostly derived from their ecological demands. Generally, not considering molecular data and cryptic species’ ecologies in monitoring programs can lead to strongly biased assessment results and ultimately to wrong management plans. The latter is especially problematic as management programmes often need to focus on protecting the maximum amount of biodiversity with the least amount of monetary effort. Using molecular methods can help to identify and effectively protect species and intraspecific diversity.

## ACKNOWLEDGMENTS

We thank Nuria Bonada for help with acquiring the sampling permission for the Parque Natural Montseny and helpful discussions. The Parque Natural Montseny is thanked for giving us permission to collect samples in the park. We thank Alexander M. Weigand for helpful discussions, Lisa Poettker for help with sampling and lab work and Raúl González Pech for translating the abstract and figure captions. We are indebted to the North-Rhine Westphalian Academy of Sciences for financial support.

## DATA ACCESSABILITY

CO1 DNA sequences:

GenBank accession numbers for *Ancylus* Clade 1: XXX-XXX. Will be available upon publication and numbers will be added accordingly.

GenBank accession numbers for *Ancylus* Clade 4: XXX-XXX. Will be available upon publication and numbers will be added accordingly.

